# Box H/ACA snoRNP regulates lipid storage through insulin signaling pathway in *Drosophila melanogaster*

**DOI:** 10.64898/2026.03.30.715344

**Authors:** Hui yang, Liu Zhao, Xuying Zhou, Xia Li, Xun Huang, Yuan Tian

## Abstract

Lipid homeostasis is essential for organismal physiology, and its disruption contributes to metabolic disorders. Using an unbiased genetic modifier screen in *Drosophila*, we identified GAR1, a core component of the box H/ACA small nucleolar ribonucleoprotein complex, as a pivotal regulator of systemic lipid storage. We show that the H/ACA snoRNP complex is essential for maintaining lipid droplet morphology in adipose tissue and preventing ectopic fat accumulation. Moreover, null mutants of *Gar1* or *Dkc1* exhibit severe developmental defects, including reduced body size and larval lethality. RNA-seq analysis revealed that *Gar1* dysfunction triggered widespread alternative splicing defects, specifically targeting key transcripts within the insulin signaling cascade, including *chico*, *Pi3K92E*, *sgg*, and *Lip4*. Furthermore, knockdown of *Gar1* impaired insulin signaling, as evidenced by the reduced membrane localization of the tGPH fluorescence. Genetic epistasis further positions GAR1 upstream of the *lin-28*/*foxo* axis, as knocking down *lin-28* or *foxo* fully rescues the lipometabolic defects in GAR1-deficient animals. These findings reveal a previously unrecognized link between the snoRNP machinery and metabolic process, establishing the box H/ACA complex as an important coordinator that integrates RNA processing with insulin-mediated nutrient sensing to ensure developmental and lipid homeostasis.

**Article summary:** Lipid metabolism is tightly controlled by multiple factors. To find new regulators, the authors performed a genetic screen and identified a small nucleolar protein GAR1 participate in fat storage and larval development. They demonstrated a critical role of box H/ACA snoRNP complex in modulating alternative splicing and balancing insulin cascade. Blocking two insulin-related genes reversed the lipid defects caused by *Gar1* loss. These findings revealed the box H/ACA complex integrates RNA processing with insulin-mediated nutrient sensing to ensure developmental and lipid homeostasis, offering a perspective for understanding the metabolic regulation network.

## Introduction

Lipids serve as the primary energy source in living organisms, and their metabolism significantly influences growth, reproduction, vitality, and disease progression (Kamareddine et al., 2018; Lim et al., 2014; Logan-Garbisch et al., 2014). Lipid droplets (LDs) are organelles specialized in storing neutral lipids, including triglycerides, sterol esters, and esterified ceramides (Chen et al., 2020; Xu et al., 2018). Pathologically, LDs accumulate in non-adipose tissues such as the liver, heart, kidney, and muscle, a condition termed ectopic fat accumulation (EFA) (Yan et al., 2017). EFA can initiate inflammatory responses that contribute to insulin resistance (Trouwborst et al., 2018), thereby elevating the risk of metabolic syndromes, including cardiovascular diseases, fatty liver disease, and hyperglycemia (Camporez et al., 2015; Ferrara et al., 2019). Moreover, excessive free fatty acids in non-adipose cells stimulate the production of reactive oxygen species (ROS) while impairing mitochondrial respiration. This process also induces endoplasmic reticulum (ER) stress responses and increases apoptosis (Ljubkovic et al., 2019; Ly et al., 2017).

Previous studies have highlighted the crucial roles of insulin and mTOR pathways in maintaining metabolic homeostasis (Caron et al., 2015; Krycer et al., 2020). In humans and other mammals, insulin is produced by pancreatic beta cells, which modulate blood glucose levels, promote cellular proliferation, and stimulate lipid storage (Lin and Smagghe, 2019). In *Drosophila*, eight insulin-like peptides (ILP1-ILP8) are secreted during nutrient deprivation and bind to the insulin receptor on target cells (Nassel et al., 2015), leading to the phosphorylation of Chico (the IRS homolog). An active Chico recruits and phosphorylates PI3K. Phosphorylated PI3K converts PIP_2_ to PIP_3_, facilitating AKT recruitment to the plasma membrane where its activity is modulated by PTEN (Zhang and Zhang, 2019). Activated AKT phosphorylates the transcription factor FOXO, preventing its nuclear translocation, and thereby inhibiting the expression of target genes such as *Lip4* and *4EBP*. The RNA-binding protein LIN-28, which is evolutionarily conserved from *C. elegans* to mammals, regulates the early developmental cell lineage formation (Ambros and Horvitz, 1984). *LIN-28A* enhances glucose uptake by augmenting insulin-PI3K-mTOR signaling, primarily by derepressing key *let-7* targets in the pathway, including IGF1R, INSR, IRS2, PIK3IP1, AKT2, TSC1, and RICTOR (Zhu et al., 2011). In *Drosophila*, *lin-28* deletion mutants exhibit reduced cell size and accelerated larval-pupal transition, whereas its overexpression delays pupation and causes lethality (Gonzalez-Itier et al., 2018).

Emerging evidence underscores the importance of non-coding RNAs and their binding proteins in cellular metabolic homeostasis. Small nucleolar RNAs (snoRNAs) range from 60 to 300 nucleotides in length and reside primarily in the nucleolus. They are categorized into three groups based on structure and protein associations: box C/D snoRNAs (snoRD), box H/ACA snoRNAs (snoRA) and orphan snoRNAs (Ojha et al., 2020; Światowy and Jagodzińśki, 2018). SnoRNAs regulates various post-transcriptional processes, including rRNA acetylation, tRNA methylation, rRNA pseudo-uridylation, and mRNA alternative splicing (Wang et al., 2025). Alternative splicing (AS) generates distinct mRNA isoforms in a tissue-specific manner, driving developmental programs, specifying tissue differentiation, and sustaining metabolic homeostasis (Einson et al., 2023; Finegan et al., 2025; Kaminska, 2025; Kanno et al., 2023; Zhang et al., 2025). For instance, the box C/D snoRNA SNORD88B has been shown to recruit the splicing factors SRSF1 and U2AF1 to regulate the alternative splicing of pre-G3BP1 (Lu et al., 2025).

Similar to box C/D snoRNAs, box H/ACA snoRNAs function as small nucleolar ribonucleoprotein (snoRNP) complexes. These complexes comprise core protein components such as DKC1 (dyskerin), NOP10, GAR1, NHP2, and NAF1 (Fatica et al., 2002; McMahon et al., 2015; Watkins et al., 1998). NAF1 is present in nascent snoRNPs, whereas GAR1 is exclusively present in mature complexes (Massenet et al., 2017). Mutations in *Dkc1*, *Nop10* and *Nhp2* are associated with human dyskeratosis congenita (Heiss et al., 1998; Vulliamy et al., 2008; Walne et al., 2007). *Dkc1* depletion induces cytoskeletal remodeling in human tumor cells (Di Maio et al., 2017) and promotes intestinal stem cell regeneration in *Drosophila* (Vicidomini et al., 2017). Meanwhile, downregulation of *Naf1, Nop10, Dkc1, and Gar1* disrupts cyst formation in *Drosophila* ovaries (Breznak et al., 2023; Morita et al., 2018). Despite documented roles of snoRNA-binding proteins in diverse biological processes, their specific functions in lipid homeostasis remain largely unknown.

*Drosophila melanogaster* is an excellent model for studying obesity, disease, aging, and development (Heier and Kuhnlein, 2018; Liu and Huang, 2013; Musselman and Kuhnlein, 2018). The previous work demonstrated ectopic lipid accumulation in *dSeipin* null mutants and flies overexpressing *DGAT* (*midway*), with double mutants showing a pronounced synergistic effect. To further explore the regulatory network of lipid metabolism, we conducted a genetic modifier screen in *Drosophila* to identify the genes involved in lipid droplet deposition and morphology. In this study, we identified GAR1, a snoRNA-binding protein, as a critical regulator of lipid homeostasis. Subsequent cellular biological analyses, RNA-seq, and genetic interaction assays revealed a functional link between box H/ACA snoRNA-binding proteins and the insulin signaling pathway in the regulation of lipid metabolism and development.

## Materials and methods

### Fly stocks and husbandry

All the fly stocks were reared on standard corn meal food. For the *DGAT* modifier screen, the *EP* strains were provided by Dr. Jiahuai Han, and the *RNAi* strains were from the NIG stock center. For the genetic interaction with genes of insulin pathway, all the *RNAi* strains were ordered from Tsinghua fly center and BDSC. The other strains used in this study were listed as following: *ppl-Gal4* (Dr. Pierre Leopold), *UAS-DGAT* (BDSC, 20167)*, Gar1* (BDSC, 34013; BDSC, 21775), *Dkc1* (VDRC, 109616; VDRC, 34597; BDSC, 36595), *Nhp2* (BDSC, 51784), *Nop10* (BDSC, 55194), tGPH reporter (BDSC, 8164), *lin-28* (Tsinghua, TH01982.N), *foxo* (BDSC, 32993) and *da-Gal4* (Tsinghua).

### Genetic screen

The *ppl-Gal4* driver was crossed with *UAS-DGAT* and designated as *ppl>DGAT* in this study. The *ppl>DGAT* females were then mated with males carrying either *EP* or *RNAi* strains. Flies were reared at 29°C to maximize GAL4 activity. Wandering third-instar larvae harboring both the *ppl>DGAT* and the *EP* or *RNAi* transgene were dissected, mounted with PBS and visualized using differential interference contrast (DIC) microscopy on a Zeiss microscope.

The list of genes obtained from the genetic screen was submitted to the Metascape online analysis platform (https://www.metascape.org), with the species set to *Drosophila melanogaster*. Gene Ontology (GO) enrichment analysis was performed using the express analysis mode with default parameters (*P*-value < 0.01, minimum count ≥ 3, and enrichment factor ≥ 1.5). The top 20 significantly enriched GO biological process terms were selected based on enrichment significance, and the enrichment heatmap was generated using the built-in function of Metascape (Zhou et al., 2019).

### Inverse PCR and sequencing

Thirty anesthetized flies were collected into a 1.5 mL tube and froze at - 80°C. Genomic DNA was extracted and digested by RsaI at 37°C. After digestion, the enzyme was inactivated at 65°C for 20 min, and the digested DNA was self-ligated using T4 DNA ligase to generate circular products. The ligated samples were amplified by PCR with primers LA(f).1 (GGGAATTGGGAATTCGTTAA) and LA(r).1 (TAGCGACGTGTTCACTTT GC). The purified PCR products were sequenced with primer LA(f)seq1 (CTCTCAACAAGCAAACGTGC). The resulting sequences were blasted against the FlyBase database to identify the affected genes. For additional details, refer to the Drosophila Gene Disruption Project (Bellen et al., 2011).

### Fluorescent fusion expression vector construction

The coding region of *Gar1* (GenBank accession number: PX393084) was amplified using primers GAATTCATGGGATTTGGTAAACCTCG and GGTACCCTACCACCGACCCCGACC, and subsequently cloned into the pEasy-T1 vector (TransGen Biotech). The resulting T1-*Gar1* plasmid was then double-digested with EcoRI-KpnI and ligated into the pUAST-attB vector. To construct *UAS-Gar1-eGFP*, the *Gar1* fragment was amplified with 4038-5BglII (AGATCTATGGGATTTGGTAAACCTCG) and 4038-3nsBamHI (GGATCCCCACCGACCCCGACCACC), digestion with BglII-BamHI, and inserted into the eGFP-N3 vector. The *Gar1-eGFP* fusion fragment was finally excised via BglII and XbaI digestion and ligated into the pUAST-attB vector.

Similarly, the coding regions of other box snoRNP components were amplified as follows: *Dkc1-RA* using primers GAATTCTCTCTCCGTCTATT-AGTTGATT and AGATCTTATTCCTGAGCTTCGTCTC; *Nhp2* using primers GAATTCATGGGCAAAGTGAAAGTAG and GGTACCGACGGGTATGTTT-AGTGC; and *Nop10* using primers CCTCGAGATGTATCTGATGTACACAA and GGTACCGTAAATGGGCTCCGGCT.

### Lipid droplet staining

To analyze lipid droplet (LD) morphology, wandering third-instar larval fat body and salivary gland were dissected, stained with BODIPY 493/503 (Invitrogen; 1 mg/mL, 1:500) or Nile Red (Sigma; 0.5 mg/mL, 1:500) together with DAPI (Sigma; 1 mg/mL, 1:10,000), and imaged using a Leica TCS SP8 confocal microscope. The sizes of lipid droplets were quantified using Image-Pro Plus 6.0 within a 150 µm * 150 µm rectangle from three images per genotype. The 30 largest lipid droplets (LDs) were used to generate violin plots. Statistical significance was determined based on mean values using GraphPad Prism 10.

### tGPH analysis

For the tGPH assay, embryos from *da-Gal4* and *RNAi* mutants were collected within a 4-hour period and reared at 29°C. Fat body and salivary gland from 68-72 hours larvae were dissected and examined under a Zeiss Axio Imager M2 microscope. In fat body cells, the fluorescent intensity was measured within 100 μm - 100 μm rectangles from 3-6 figures for each genotype. The cellular segmentation was performed using the online version of Cellpose 4.0. Following this, membrane and cytosolic signals were measured separately using ImageJ (Fiji). The resulting data were plotted and analyzed with GraphPad Prism 10.

For gray value measurement in the salivary gland, the intensity was measured within 50 μm - 50 μm rectangles from five figures for each genotype. A line was drawn between the nuclei of two adjacent cells to quantify the fluorescence intensity distribution along this line using ImageJ (Fiji). For the distribution curve, three datasets from different cells per genotype were selected for normalization of both the X-axis (distance) and Y-axis (gray value). Subsequently, the normalized fluorescence distribution profiles were subjected to curve fitting, generating a lowess fit curve for each genotype.

### Knockout strategy and larval morphological analysis

Two sgRNAs were designed following an established online protocol (Housden et al., 2015; https://www.flyrnai.org/crispr3/web/) and inserted into the pUAST-attB vector using Golden-Gate cloning. The resulting sgRNA-expressing flies were subsequently crossed with *Act-Cas9* flies. The deletion mutants were identified by PCR amplification followed by sequencing. Heterozygous alleles were balanced using the CyO**·**P(ActGFP) JMR1 chromosome.

Embryos were collected within a 4-hour interval. After 24 hours, homozygous mutant larvae (lacking GFP fluorescence) were sorted under an Olympus SZX16 stereomicroscope. Morphological observation was carried out using Olympus BX53F or Motic K-400L microscopes. Body length measurements were obtained with Motic Images Advanced 3.2 software, and the resulting data were summarized in column graphs and subjected to statistical analysis using GraphPad Prism 10.

### Transcriptome analysis

Embryos from the *w^1118^* and *Gar1^2^* mutants were collected within 4 hours. After 24 hours, wild-type or non-GFP mutant larvae (homozygotes) were transferred to fresh dishes, and rinsed with PBS in order to eliminate contaminants. After drying on filter paper, larvae were immersed in 1mL of Trizol solution (B511311, Sangon, China) for RNA stabilization and extraction. RNA sequencing was performed by Sangon Biotech Co., Ltd. (Shanghai, China). RNA quality was assessed using a NanoPhotometer® spectrophotometer (IMPLEN, CA, USA) and a Qubit® 2.0 Fluorometer (Invitrogen). Sequencing libraries were constructed using the VAHTSTM mRNA-seq V2 Library Prep Kit for Illumina® with 1 μg of total RNA as input. After sequencing, raw data quality was evaluated using FastQC (version 0.11.2), and reads were trimmed using Trimmomatic (version 0.36). Clean reads were mapped to the reference genome using HISAT2 (version 2.0) with default parameters. The differential gene expression analysis was performed using DESeq2 (version 1.12.4), with genes satisfying *q*-value < 0.05 and |FoldChange| ≥ 2 considered significantly differentially expressed. Alternative splicing events were analyzed using rMATS (Shen et al., 2014), and significant events were filtered with FDR < 0.05 and |IncLevelDifference| > 0.1.

### Q-PCR analysis

The late L3 larvae were collected, washed with PBS in order to remove any impurities, dried on filter paper carefully, and subsequently mounted in 1mL of Trizol solution (Invitrogen). The RNAs were extracted by Trizol-chloroform methods. The first strand cDNAs were obtain using oligo dT primer and M-MLV Reverse Transcriptase (Invitrogen). The specific primers for each gene were designed from online database in NCBI. The real-time quantitative polymerase chain reactions used SYBR green I (ABI). The column chart and statistical analysis were generated using GraphPad Prism 10.

## Results

### A modifier screen for genes involved in lipid storage

To identify the genes regulating lipid metabolism, we conducted a genetic modifier screen in *Drosophila melanogaster* by co-expressing *DGAT*, which encodes a diacylglycerol acyltransferase (Chen et al., 2022). Compared to the wild type, overexpression of *DGAT* led to numerous small puncta in the salivary glands (black arrows, Fig. 1a and 1a’) and large droplets in the fat bodies of third-instar larvae (Fig. 1g). To investigate the genetic regulators of lipid metabolism, we crossed *DGAT*-overexpressing flies (*ppl>DGAT*) with *EP* or *RNAi* strains. Genes in *RNAi* strains were obtained from an online database, whereas those in *EP* strains were identified using inverse PCR (Fig. S1a and S1b; Table 1). We then dissected the fat body and salivary gland tissues from third-instar larval offspring and analyzed LD’ morphology using DIC microscopy. From a total of 2100 EP and 1800 RNAi strains screened, we identified 109 genes implicated in the regulation of lipid metabolism (Table 1). GO enrichment analysis clustered these genes into diverse biological processes, primarily glycerophospholipid metabolism, triglyceride biosynthesis, neural development, chromatin organization, homeostatic process, and reproduction (Fig. S1c). Based on the morphology changes in the lipid droplets, we categorized the observed phenotypes into seven distinct classes (Table 1). Classes I-V exhibited prominent differences in the salivary glands, whereas classes VI and VII showed alterations primarily in the fat body.

**Fig. 1.**
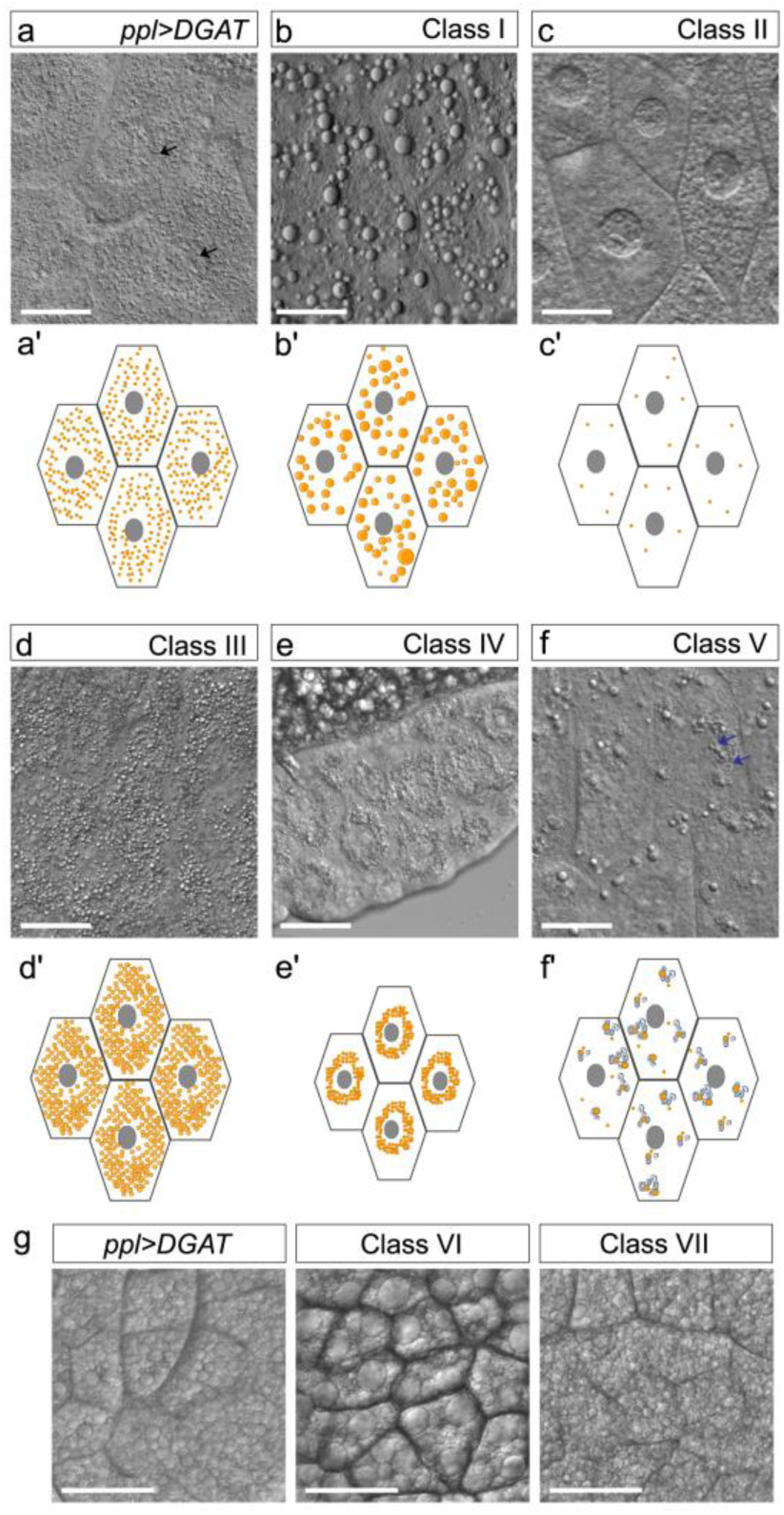
The double mutants based on *ppl>DGAT* exhibit seven distinct morphological types of lipid droplets. a, a’) *DGAT* overexpression induced ectopic lipid droplet accumulation. b–f) The changes in ectopic fat accumulation in larval salivary gland. b’–b’) The schematic diagrams of b–f, respectively. g) The changes of lipid droplets in larval fat body, and *ppl>DGAT* was used as a control. Scale bar: 50 μm. **ALT TEXT :** Graphs depict seven different patterns of lipid deposition changes isolated from the genetic screen based on overexpression of a triglyceride synthesis enzyme, with subfigures labelled from a to g. b to f represents prominent differences in the salivary glands, which is enhanced or diminished lipid accumulation in the salivary gland. g shows alterations increased or decreased sizes of lipid droplets in the fat body.

**Table 1.**
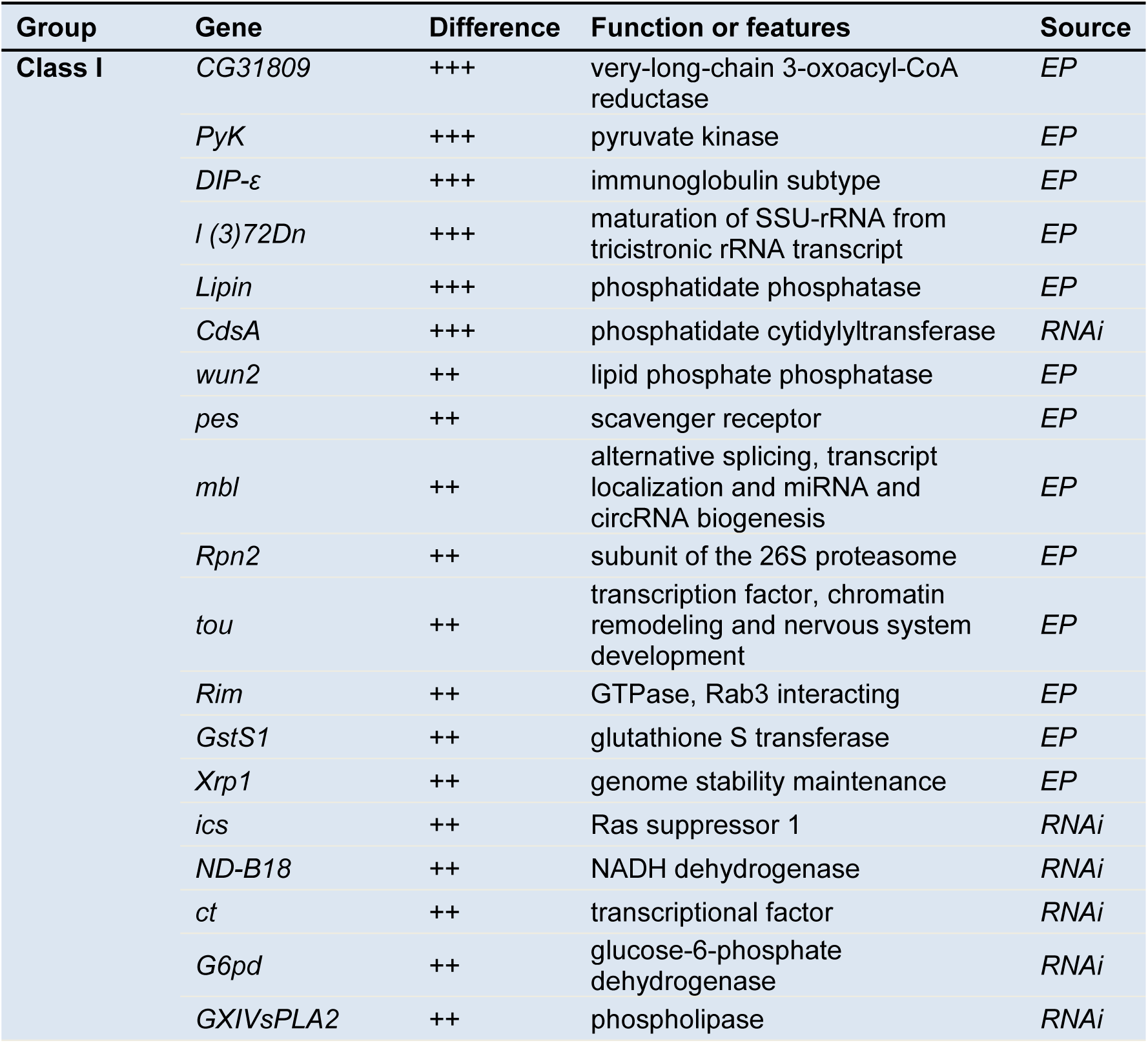
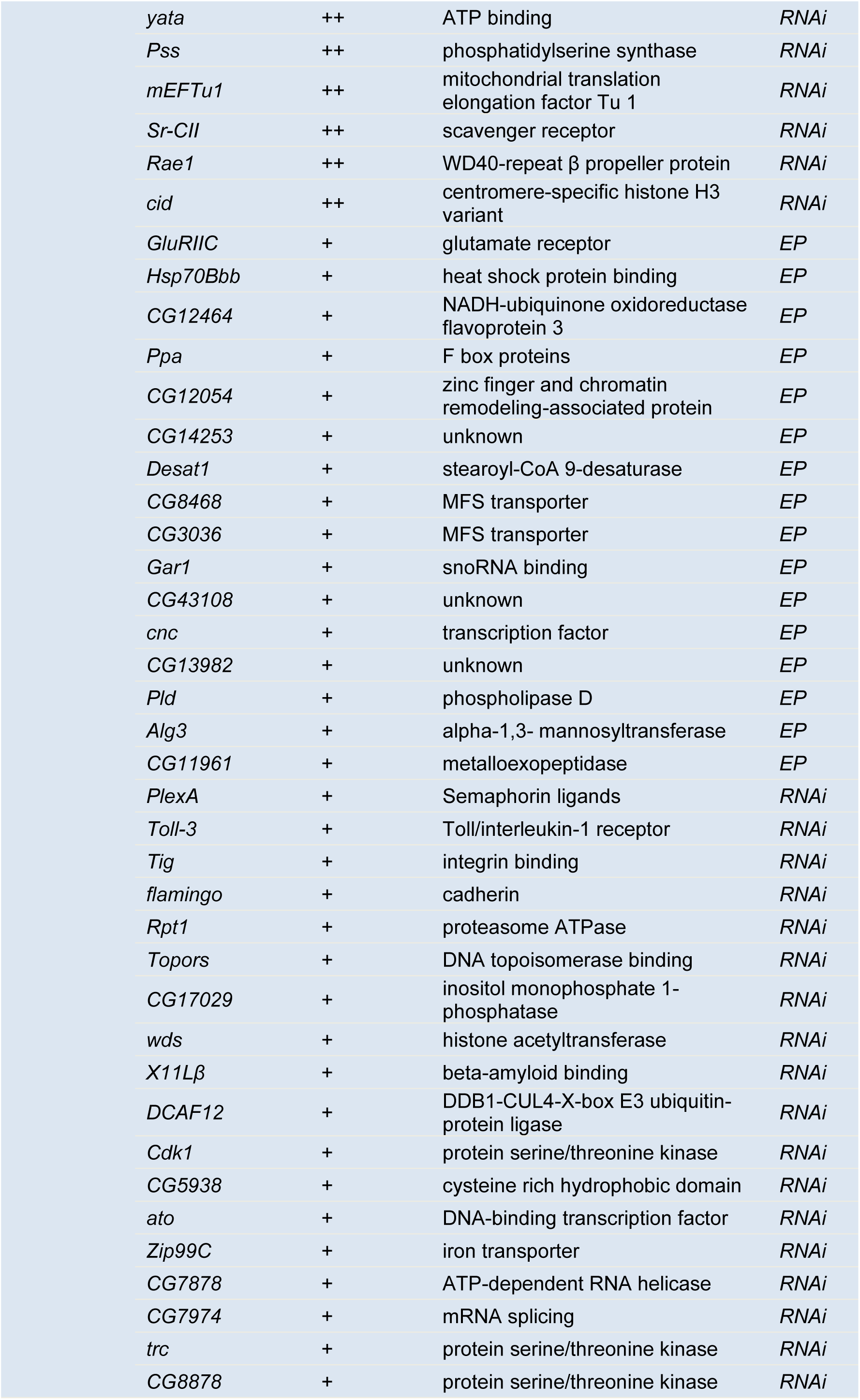
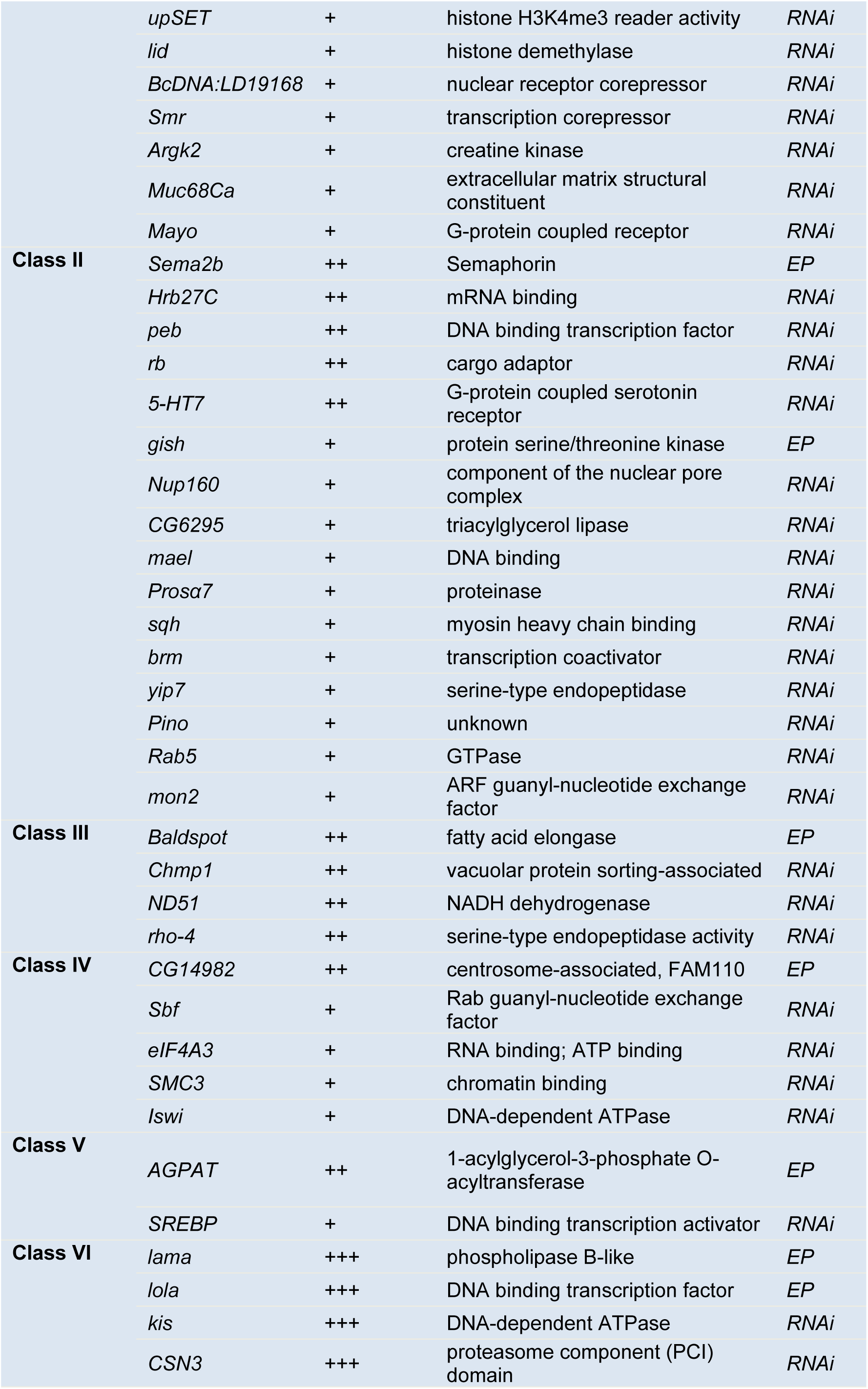
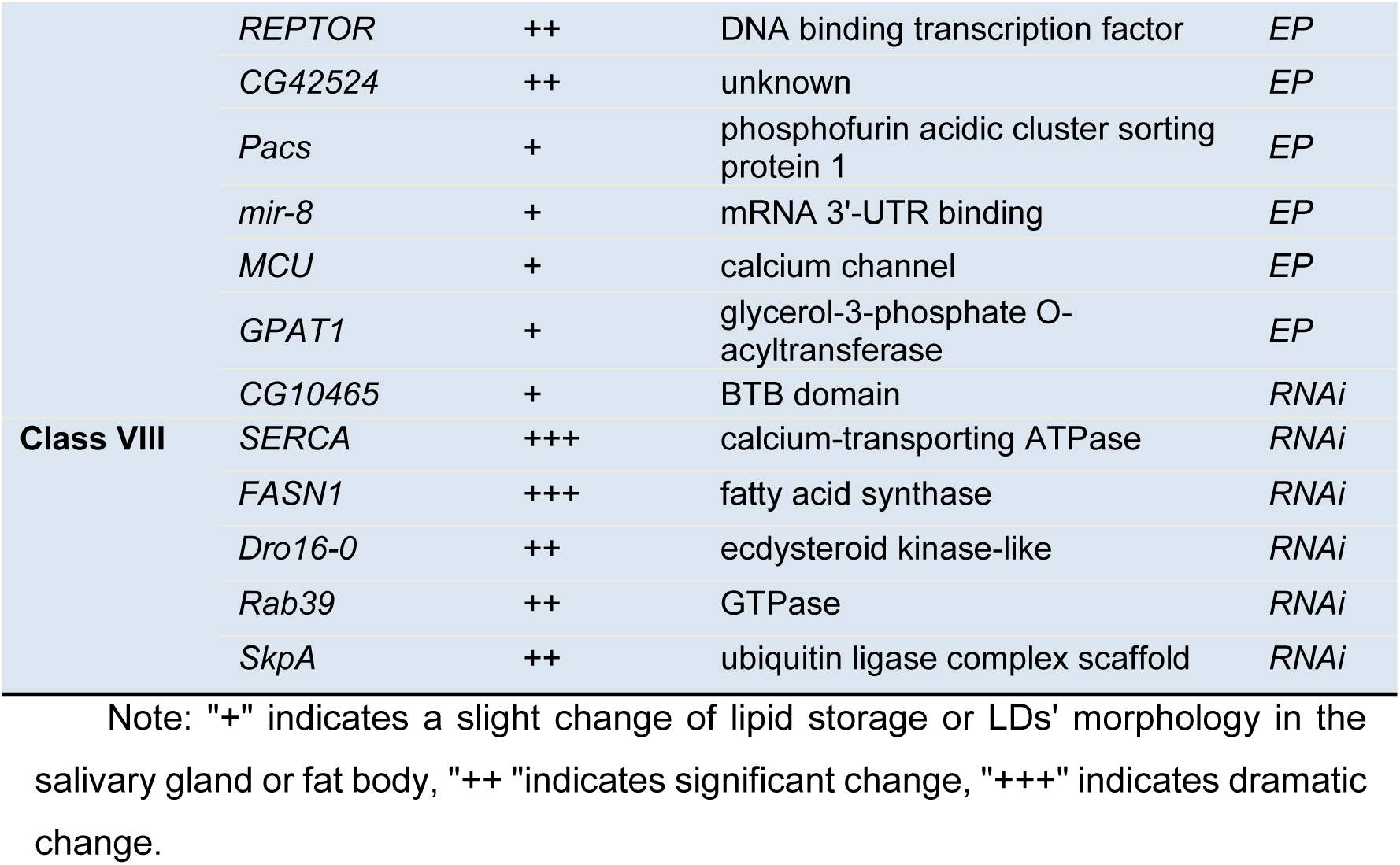
Genes related to ectopic lipid accumulation identified in the screen.

The class I double mutants displayed pronounced accumulation of supersized lipid droplets in the salivary gland, resembling those typically found in the fat body (Fig. 1b and 1b’). This class includes several well-characterized lipid metabolism related genes such as *Lipin*, *wun2*, *CdsA*, *Desat1,* and *GluRIIC* (Table 1). In contrast, class II mutants showed reduced lipid accumulation in the salivary gland, with few or no cytoplasmic puncta (Fig. 1c and 1c’). Genes in this class include *Rab5*, *Sema2b,* and *Nup160* (Table 1). Notably, class III double mutants displayed exhibited substantial accumulation of numerous small lipid droplets in salivary gland, along with a moderate increase in lipid droplet size (Fig. 1d and 1d’). This category included genes encoding fatty acid elongase (*Baldspot*) and calcium ion-binding protein (*rho-4*) (Table 1). Interestingly, class IV was characterized by the presence of multiple small cytoplasmic lipid droplets clustered around and enclosing the nucleus, in contrast with the dispersed pattern observed in the control group (*ppl>DGAT*). This phenotype was often accompanied by reduced salivary gland cell size (Fig. 1e and 1e’). Associated genes included *SMC3*, *Iswi,* and *Sbf* (Table 1). Moreover, the class V mutants featured large cytoplasmic lipid droplets surrounded by membrane-like circular structures (blue arrows in Fig. 1f and 1f’). Notably, this group included the lipogenic genes, *AGPAT* and *SREBP* (Table 1).

In contrast to the diverse salivary gland phenotypes, changes in the fat body were characterized by alterations in lipid droplet size, with droplets showing either enlargement or reduction. Class VI mutants exhibited enlarged lipid droplets in the fat body and contained genes encoding *Lama*, *Kis*, and *Lola* (Fig. 1g; Table 1). Class VII mutants, on the other hand, displayed reduced lipid droplet size in the fat body and involved genes such as *SERCA*, *rab39*, and *FASN1* (Fig. 1g; Table 1). In summary, this screen uncovered critical genes involved in ectopic fat accumulation, prompting further investigation into their roles in the regulation of lipid metabolism.

### Subcellular Localization of box H/ACA snoRNP

From this screen, we found that overexpression of *Gar1* and *DGAT* simultaneously led to an increase in EFA in the salivary gland (Fig. S2a; Table 1). GAR1 is a snoRNA binding protein that associates with other proteins and box H/ACA snoRNAs to form the snoRNP complex, thereby regulating multiple cellular processes (Breznak et al., 2023). However, the functional role of the snoRNP complex in lipid metabolism remains unclear. Fluorescence imaging was performed to systematically examine the subcellular localization patterns of these proteins. We amplified and sequenced the coding sequences of *Gar1* and found that the cloned sequence contained a 6-nucleotide deletion relative to the FlyBase reference, which resulted in a polypeptide lacking two glycine residues (Fig. S2b). To determine whether cloned GAR1 was functional, we expressed UAS-GAR1-eGFP using the fat body-specific driver *ppl-Gal4*. As expected, GAR1-eGFP was exclusively localized to the nucleus and exhibited a punctate distribution in both the larval salivary glands and fat body cells (Fig. 2a). This localization pattern was confirmed by co-staining with phalloidin and DAPI (for nuclei), and is consistent with previous reports in mammalian systems (Chamousset et al., 2010; Jacob et al., 2013). Furthermore, *da-Gal4*-driven overexpression of GAR1 in the *Gar1^2^* mutant background fully rescued the mutant phenotype to that of the wild type, demonstrating that GAR1 was successfully cloned (Fig. S2c, S2d, 4k). Based on these findings, we extended analysis to other core H/ACA snoRNPs using similar fluorescent tagging strategies. The sequences of *Dkc1-RA*, *Nop10*, and *Nhp2* were 100% identical to the reference sequences. Interestingly, DKC1, NHP2, and NOP10 exhibited similar localization, with predominant nuclear localization and faint cytoplasmic signals in fat body cells (Fig. 2b and 2c). These patterns resembled the localization of CBF5-eYFP (DKC1-eYFP) in *Arabidopsis thaliana* (Lermontova et al., 2007). The above findings demonstrate that the four core proteins exhibit conserved localization patterns. Coupled with the initial observation that *Gar1* overexpression promotes ectopic lipid accumulation, it suggests that the entire complex plays a critical role in regulating specific physiological processes, such as lipid homeostasis. Nevertheless, the integrated function of the snoRNPs in lipid metabolism remains to be determined, prompting us to investigate their role in *Drosophila*.

**Fig. 2.**
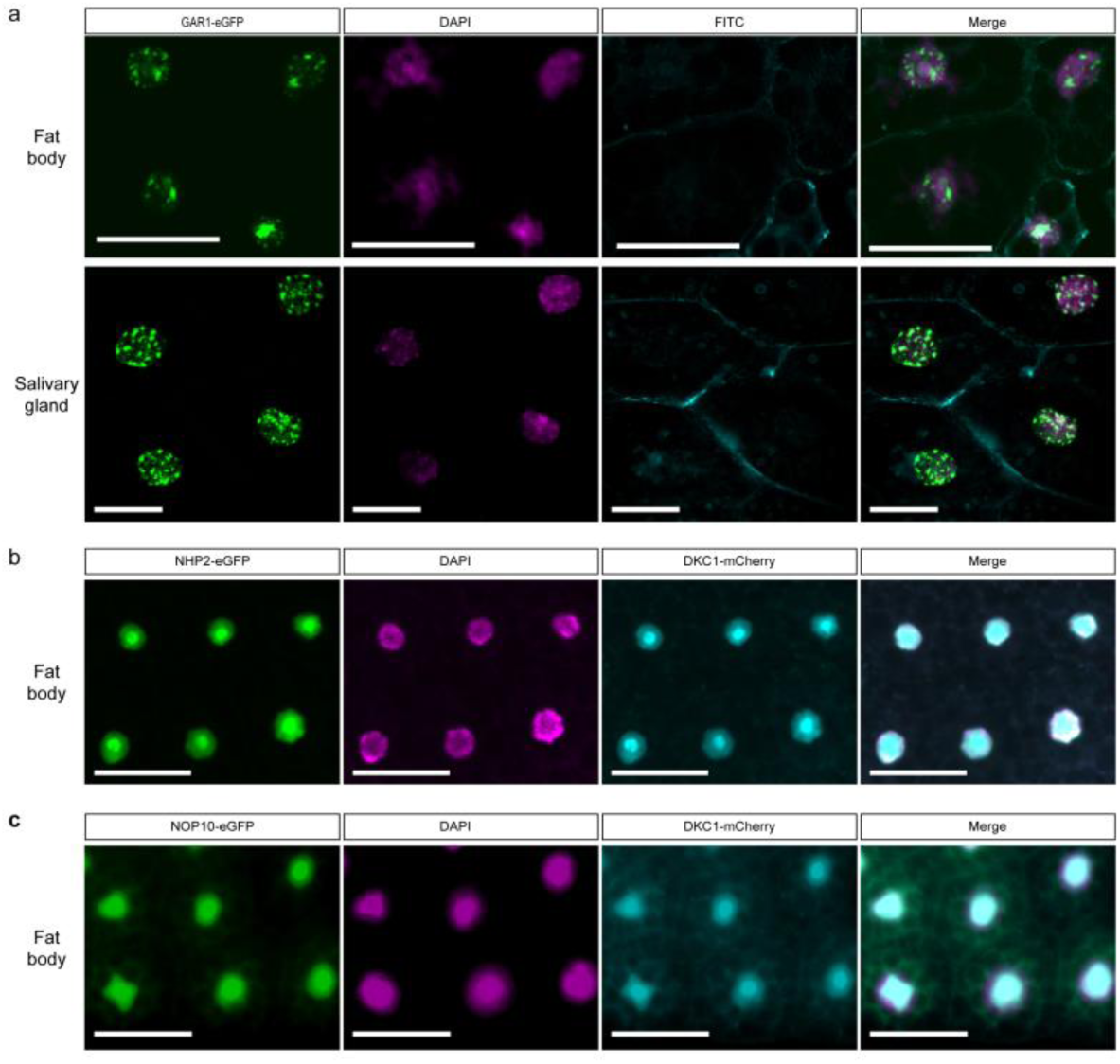
The localization of box H/ACA snoRNA binding proteins. a) GAR1-eGFP fusion proteins showed punctate distributions in the nuclei of the salivary gland and fat body. The top panel: fat body, the bottom panel: salivary gland. Green (GAR1-eGFP), Cyan (Phalloidin-FITC), Magenta (DAPI). b) DKC1 was co-localized with NHP2. Cyan (DKC1-mCherry), Green (NHP2-eGFP), Magenta (DAPI). c) DKC1 was co-localized with Nop10. Cyan (DKC1-mCherry), Green (Nop10-eGFP), Magenta (DAPI). Scale bar: 50 μm. **ALT TEXT :** Graphs illustrate the subcellular localization of four core box H/ACA snoRNA binding proteins. a) shows GAR1 localized in the nucleus with a punctate pattern. (b) and (c) depict DKC1 highly expressed in the nucleus without puncta, colocalized with NOP10 and NHP2.

### Box H/ACA snoRNP regulates lipid homeostasis

To investigate their function *in vivo*, we analyzed changes in lipid deposition in the corresponding knockdown and overexpression flies. Salivary glands were dissected at the wandering third-instar larval stage and stained with BODIPY 493/503. We found that knockdown of *Gar1* or *Dkc1* reduced the salivary gland cell size and induced numerous small ectopic lipid droplets in the salivary glands (Fig. 3a, S2a and S3a). In contrast, double mutants combining *Gar1* or *Dkc1* RNAi with *DGAT* overexpression showed dramatically reduced salivary gland cell size, along with a mild increase in ectopic lipid accumulation (Fig. 3a). These findings suggest that both *Gar1* and *Dkc1* are crucial for suppressing EFA formation in the salivary glands. To further explore their role in EFA formation, we overexpressed *Gar1* and found that it alone did not produce a significant phenotype compared with the wild type (Fig. 3b). However, co-overexpression of *Gar1* and *DGAT* triggered the formation of large ectopic lipid droplets (Fig. 3b and S2a), which was consistent with the results of previous genetic screen. This indicates that while *Gar1* overexpression alone does not induce EFA, it strongly enhances EFA formation when *DGAT* activity is elevated. Collectively, these results demonstrate that box H/ACA snoRNP components act as key regulators of ectopic lipid accumulation in *Drosophila melanogaster*.

**Fig. 3.**
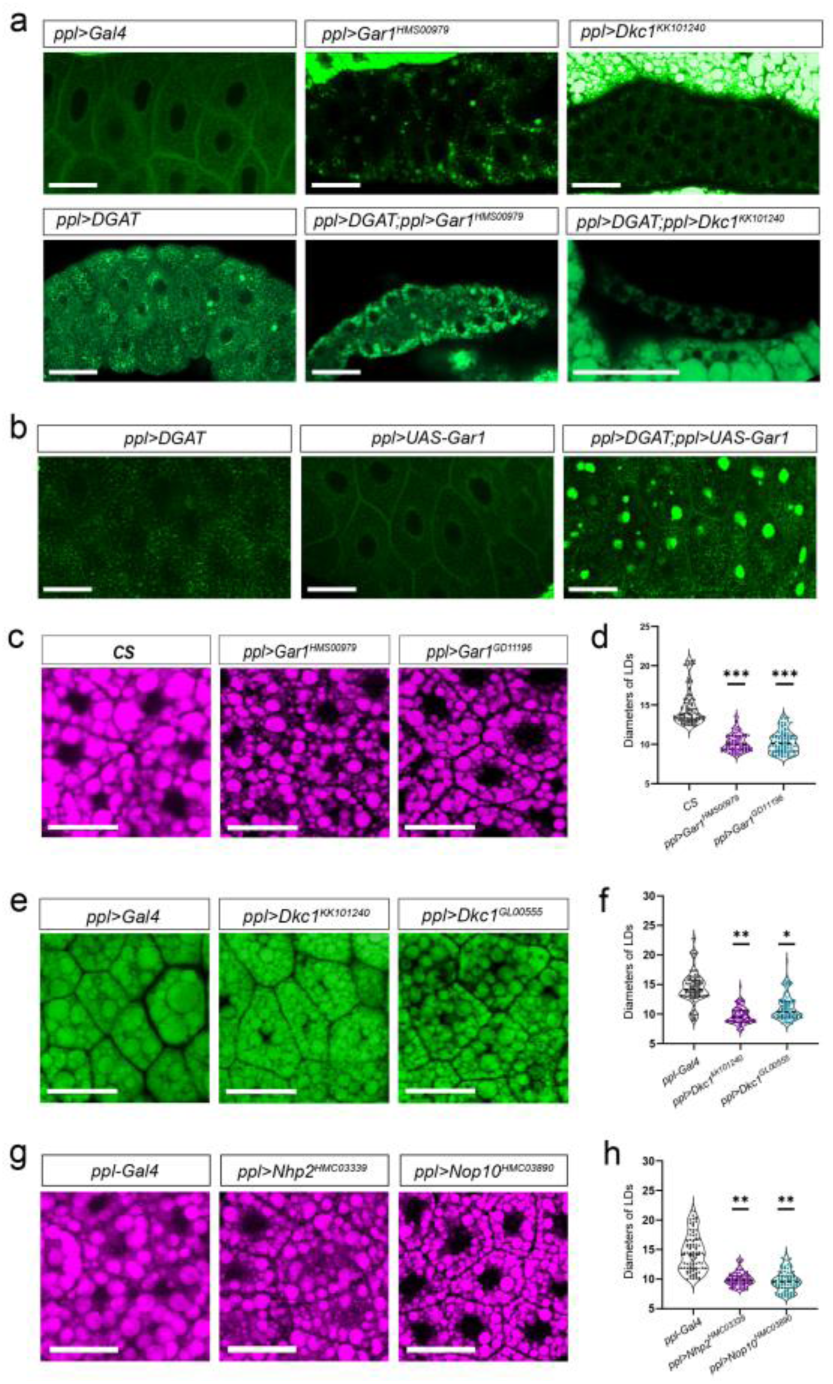
Dysfunction of *Gar1* or *Dkc1* exhibited ectopic lipid accumulation in the salivary gland and reduced lipid droplet sized in the fat body. a) Knockdown of *Gar1* or *Dkc1* led to ectopic lipid deposition. b) Overexpression of *Gar1* enhanced ectopic fat accumulation in the salivary gland at the wandering stage. c) Knockdown of *Gar1* by *ppl-Gal4* resulted in smaller lipid droplets in the fat body at the wandering stage. e, g) Reduced lipid droplet sizes were also observed in knockdown of other snoRNP-encoding genes. Genotypes are indicated in the figure. All RNAi lines were driven by UAS-GAL4 system, and the ‘*ppl>*’ was an abbreviation for *ppl>Gal4*. Lipid droplets were stained with Bodipy 493/503 (a, b, e) or Nile red (c, g). Scale bar: 50 μm. d, f, h) LD’ diameters in control and mutants. Violin plots show Max30 LDs from three figures in each genotype (n = 90 LDs per genotype). Box plots inside violins show median and quartiles. Statistical analysis was determined by one-way ANOVA using mean values, followed by Dunnett’s multiple comparisons test. d) The LD’ diameters in control and *Gar1* mutants (*CS* vs. *ppl>Gar1^HMS00979^*, df = 6, adjusted *p*=0.0003; *CS* vs. *ppl>Gar1^GD11196^*, df = 6, adjusted *p* = 0.0004). f) The LD’ diameters in control and *Dkc1* mutants (*ppl-Gal4* vs *ppl>Dkc1^kk101240^*, df = 6, adjusted *p* = 0.0075; *ppl-Gal4* vs *ppl>Dkc1^GL00555^*, df = 6, adjusted *p* = 0. 0368). h) The LD’ diameters in control, *Nhp2* mutant and *Nop10* mutant (*ppl-Gal4* vs *ppl>Nhp2^HMC03339^*, df = 6, adjusted *p* = 0.0051; *ppl-Gal4* vs *ppl>Nop10^HMC03890^*, df = 6, adjusted *p* = 0.0035). ***, *p* < 0.001; **, *p* < 0.01; *, *p* < 0.05. **ALT TEXT :** Graphs depict that dysfunction of snoRNP display abnormal lipid deposition. a and b show lipid droplet staining in the salivary glandes. c), e) and g) show reduced lipid droplet size in the fat bodies following knockdown of all four box snoRNP components, with corresponding statistical analyses presented in d), f) and h), respectively.

Next, we investigated whether the dysfunction of box H/ACA snoRNP components affected lipid storage in the fat body. Lipid droplets in larval L3 fat bodies were stained with BODIPY 493/503 or Nile red and visualized using confocal microscopy. For each image, a 150 μm * 150 μm area was selected, and the diameters of the 30 largest lipid droplets (Max30 LDs) within each region were measured and compared. The average Max30 LDs’ size in *Canton-S* flies was 14.37 μm, whereas it was reduced to 10.07 μm in *ppl>Gar1^HMS00979^* flies and 10.16 μm in *ppl>Gar1^GD11196^* flies, respectively (Fig. 3c and 3d), indicating that *Gar1* knockdown decreases lipid droplet size. To determine whether *Dkc1* downregulation similarly affects lipid storage, we analyzed LD morphology in the fat bodies of flies expressing *ppl-Gal4* driven *Dkc1 RNAi* (*Dkc1^kk101240^* and *Dkc1^GL00555^*). The average Max30 LDs’ sizes were 9.75 μm and 11.13 μm, respectively (Fig. 3e and 3f), confirming that DKC1 also promotes lipid storage in the adipose tissue. We evaluated the impact of *Nhp2* and *Nop10* knockdown on lipid metabolism. As expected, *ppl-Gal4***-**driven expression of *Nhp2^HMC03339^* and *Nop10^HMC03890^* exhibited smaller fat body lipid droplets, with average Max30 LD’s sizes of 10.01 μm and 9.66 μm, respectively (Fig. 3g and 3h). Collectively, these results demonstrate that box H/ACA snoRNP dysfunction reduces lipid droplet size in the fat body and promotes ectopic lipid droplet accumulation in the salivary gland. The consistent phenotype across multiple components suggests that these four proteins function as integrated complexes to regulate lipid storage, underscoring the critical role of box H/ACA snoRNP in maintaining lipid homeostasis.

### Dysfunction of box H/ACA snoRNP component resulted in developmental defects

In addition to lipid metabolism, we explored the functions of box H/ACA snoRNPs during development. Strikingly, functional impairment of either *Gar1* or *Dkc1* results in severe developmental defects. Wild-type flies typically complete metamorphosis and eclose as adults 10 days after egg laying (AEL) at room temperature, whereas knockdown of *Gar1* (*Gar1^HMS00979^* and *Gar1^GD11196^*) driven by *ppl-Gal4* at 29°C led to pupal arrest and failed eclosion (Fig. 4a), indicating that GAR1 is essential for development. We examined whether *Dkc1* affects the developmental process. As expected, ubiquitous knockdown of *Dkc1* (*Dkc1^kk101240^*and *Dkc1^GL00555^*) using *da-Gal4* at 29°C resulted in developmental arrest at 4 days AEL and 8 days AEL, respectively (Fig. 4b and 4e). After hatching, the larvae progress through three distinct instar stages (L1, L2, and L3), each characterized by specific morphological features of the mouth hooks and tracheal system, which serve as key developmental markers (Hamid and Mishra, 2021). The L1 larvae exhibited mouth hooks with a single tooth and lacked paired anterior spiracles (Fig. S4a and S4a’). The L2 larvae developed mouth hooks bearing 2-4 teeth and possessed rounded or club-shaped anterior spiracles (Fig. S4b and S4b’). The L3 larvae displayed mouth hooks with 9-12 teeth and branched anterior spiracles (Fig. S4c and S4c’). On day 4 AEL, *da>Dkc1^kk101240^* larvae exhibited a slimmer body size. However, their overall body length was comparable to that of wild-type larvae (Fig. 4b). They showed obvious developmental abnormalities: mouth hooks possessed only three teeth (Fig. 4c), and tracheal termini were club-shaped (Fig. 4d), indicating that the mutants were arrested at the L2 stage. By 6 d AEL, wild-type larvae initiated pupariation, whereas *da>Dkc1^GL00555^* flies exhibited typical larval morphology. The mutant larvae had one tooth in the mouth hooks (characteristic of L1) and club-shaped tracheae (characteristic of L2) (Fig. 4e), suggesting developmental arrest at the L1-L2 transition. The above results indicate that knockdown of box H/ACA components leads to developmental defects.

**Fig. 4.**
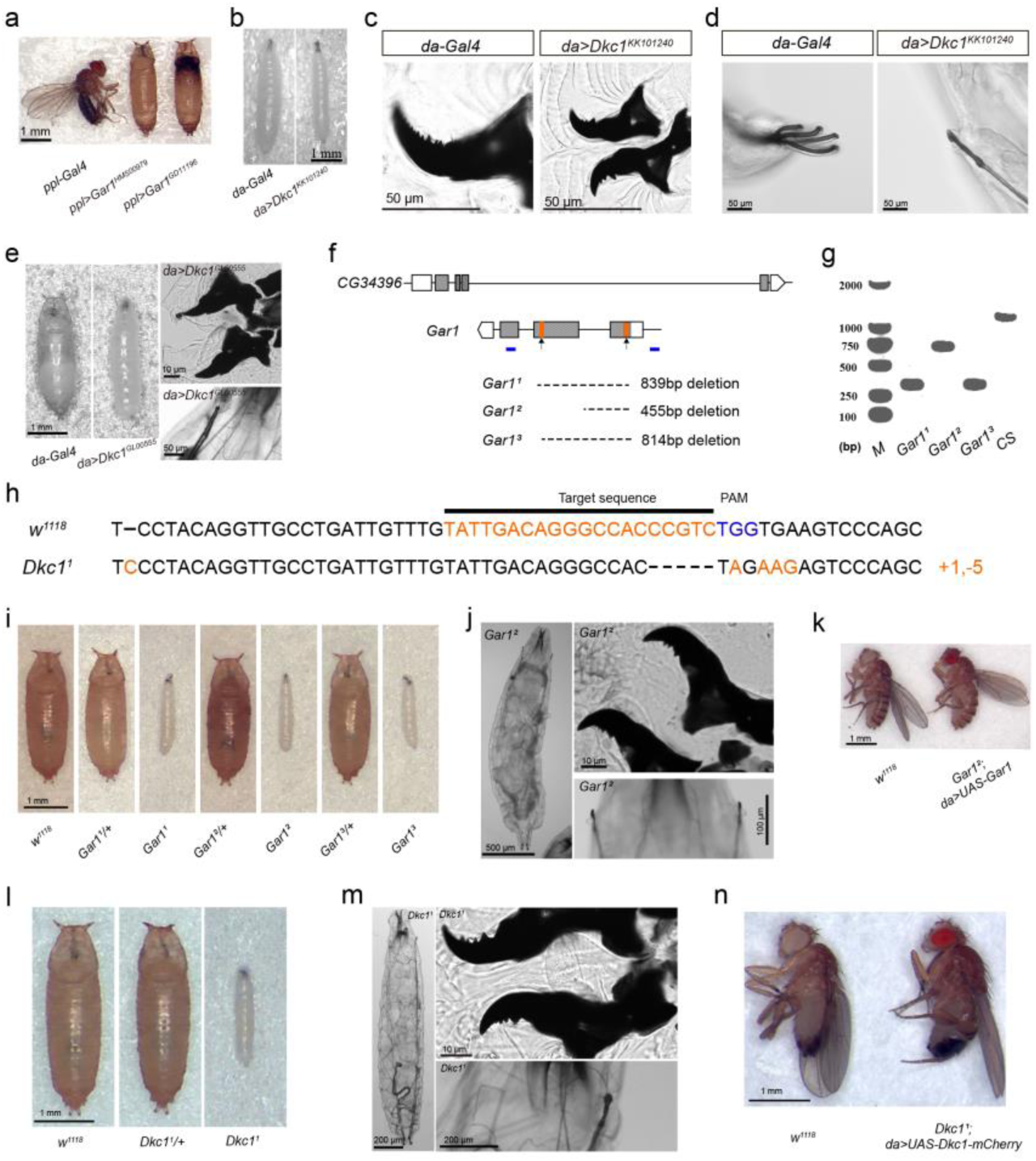
Loss of *Gar1* or *Dkc1* function causes developmental delay and arrest at the larval stage. a) Knockdown of *Gar1* results in pupal lethality when raised at 29°C driven by *ppl-Gal4*. b–e) *da-Gal4-*driven knockdown of *Dkc1* at 29°C caused larval lethality by 4 days AEL (b–d) and 6 days AEL (e). f) Genomic locus of *Gar1* and its deficiency alleles. The *Gar1* gene is located within the intron of *CG34396*. Deleted regions are indicated by dashed lines; sgRNA target sites are marked by orange rectangles. g) PCR of *Gar1* mutants and wild-type controls. Amplification was performed using primers shown as blue boxes in (f). M, DNA marker. h) Schematic diagram of *Dkc1^1^* mutant. i) *Gar1* loss of function mutants exhibited developmental arrest on 8 d AEL. j) Mouth hook and spiracle morphology of *Gar1^2^* mutant on 8 d AEL. k) The morphology of wild-type and *UAS-Gar1* rescued flies at 3 d post-eclosion, raised at 25°C. l) *Dkc1^1^* mutant exhibited developmental arrest on 8 d AEL. m) Mouth hook and spiracle morphology of *Dkc1^1^* mutant on 8 d AEL. n) The morphology of wild-type and *UAS-Dkc1-mCherry* rescued flies at 5 d post-eclosion, raised at 25°C. Scale bars were indicated in the relative figures. **ALT TEXT :** Graphs depict that dysfunction of snoRNP disrupt larval developmental process, with subfigures labelled from a to n. (a–e), (i, j) and (l, m) show *Gar1* and *Dkc1* mutants died at 1^st^ and 2^nd^ larval stage, as characterized by the morphological changes in the mouth hook and tracheal. f–h) illustrate the deletion sites in the relative mutants. k and n) illustrate the mutants could be restored to the adult stage by overexpressing the wild-type copy of GAR1 or DKC1.

To further elucidate the loss of function phenotypes, we generated deletion mutants of *Gar1* and *Dkc1* using CRISPR-Cas9. For *Gar1*, the two sgRNAs targeting the first and second exons yielded three deletion mutants (*Gar1^1^*, *Gar1^2^*, *Gar1^3^*), with deleted regions of 839 bp, 455 bp, and 814 bp, respectively (Fig. 4f and 4g). Since *Gar1* is located within an intron of *CG34396*, we assessed whether its deletion affected *CG34396* expression and found no significant difference in mRNA levels between the wild-type and *Gar1^2^* mutants (Fig. S3b). Similarly, we generated a *Dkc1* null mutant (*Dkc1^1^*) containing a 1-bp insertion and a 5-bp deletion that causes premature translational termination (Fig. 4h). Similar to the RNAi mutants, the loss of function in both *Gar1* and *Dkc1* resulted in developmental arrest and larval lethality. On day 2 AEL, the *Gar1^2^*, *Dkc1^1^* and wild type exhibited similar morphology, with one mouth hook tooth and no anterior spiracles (Fig. S5a–S5c’), indicating synchronous development. By 3 d AEL, the wild-type larvae had transitioned to the L2 stage (Fig. S6a and S6a’), whereas both L1 larvae (Fig. S6b–S6c’), and L2 larvae were present in *Gar1^2^* and *Dkc1^1^* mutants (Fig. S6d–S6e’). On day 4 AEL, wild-type larvae reached the L3 stage (with 11 teeth and branched spiracles) (Fig. S7a and S7a’), whereas mutants exhibited only three teeth and club-shaped spiracles (Fig. S7b–S7c’), confirming developmental delay. By 5 d AEL, mutant showed severe growth impairment, with body lengths of 1.3-1.8 mm compared to 3.8-4.6 mm in control animals (Fig. S8a–S8c). All mutants died between 6 and 11 d AEL, and the lethality could not be improved by 20-Hydroxyecdysone addition (Fig. S8d and S8e). By 8 d AEL, wild-type animals had proceeded to the pupal stage, whereas *Gar1^2^* and *Dkc1^1^* mutants exhibited developmental arrest at the larval stage (Fig. 4i and 4l), displaying L2 features such as club-shaped spiracles and four mouth hook teeth (Fig. 4j and 4m). Genetic rescue of *Gar1^2^* using *da-Gal4* to drive *UAS-Gar1* expression restored the normal development, viability, fertility, and body length (Fig. 4k). Similarly, *da-Gal4*-driven overexpression of *Dkc1-mcherry* rescued *Dkc1^1^* mutants from the wild-type phenotype (Fig. 4n). In summary, these results demonstrate that the box H/ACA snoRNP complex is essential for individual development in *Drosophila*.

### Box H/ACA snoRNP regulates lipid homeostasis through insulin signaling pathway

To investigate the regulatory role of *Gar1* in lipid metabolism and development, we performed the transcriptomic profiling of *Gar1* loss-of-function mutant (*Gar1^2^*). RNA was extracted from the L1 larvae of both wild-type and *Gar1^2^* mutants, and their transcriptomes were compared. This analysis identified 559 differentially expressed transcripts, with 173 upregulated and 395 downregulated in the mutant relative to the wild type (Fig. S9a). Subsequent Gene Set Enrichment Analysis (GSEA) revealed the upregulation of processes related to nucleotide stability and the concurrent downregulation of both fatty acid and lipid metabolic processes in *Gar1^2^* mutants (Fig. 5a). This downregulation is consistent with the observed reduction in lipid storage in the fat body upon *Gar1* silencing. Because the snoRNP complex has been reported to be involved in RNA splicing, we analyzed alternative changes between *Gar1^2^* homozygous mutants and wild-type animals. We detected five types of AS events: skipped exon (SE), mutually exclusive exon (MXE), alternative 5’-Splice site (A5SS), alternative 3’-Splice site (A3SS) and retained intron (RI). Notably, 366 significant differentially spliced events were identified (Fig. S9b), including A5SS (14.75%), A3SS (22.40%), MXE (6.83%), RI (22.68%), and SE (33.33%). Among these, splicing factor encoding genes displayed significant AS changes, including *Rbfox1* and *HnRNP-K* (Fig. 5b). Interestingly, we also found key components of the insulin signal showed AS changes in the *Gar1^2^* mutant flies, including *chico*, *Pi3K92E* (*dp110*), *sgg* (*GSK-3β*), and *Lip4* s (Fig. 5b–5e). Notably, AKHR, an antagonist of insulin signaling, also exhibited AS changes (Fig. 5b). Meanwhile *SREBP*, which encodes a sterol regulatory element-binding protein, underwent alternative splicing in an A3SS manner (Fig. 5b and 5f) These splicing alterations in the insulin pathway genes suggest the potential disruption of insulin signaling in *Gar1* mutant.

**Fig. 5.**
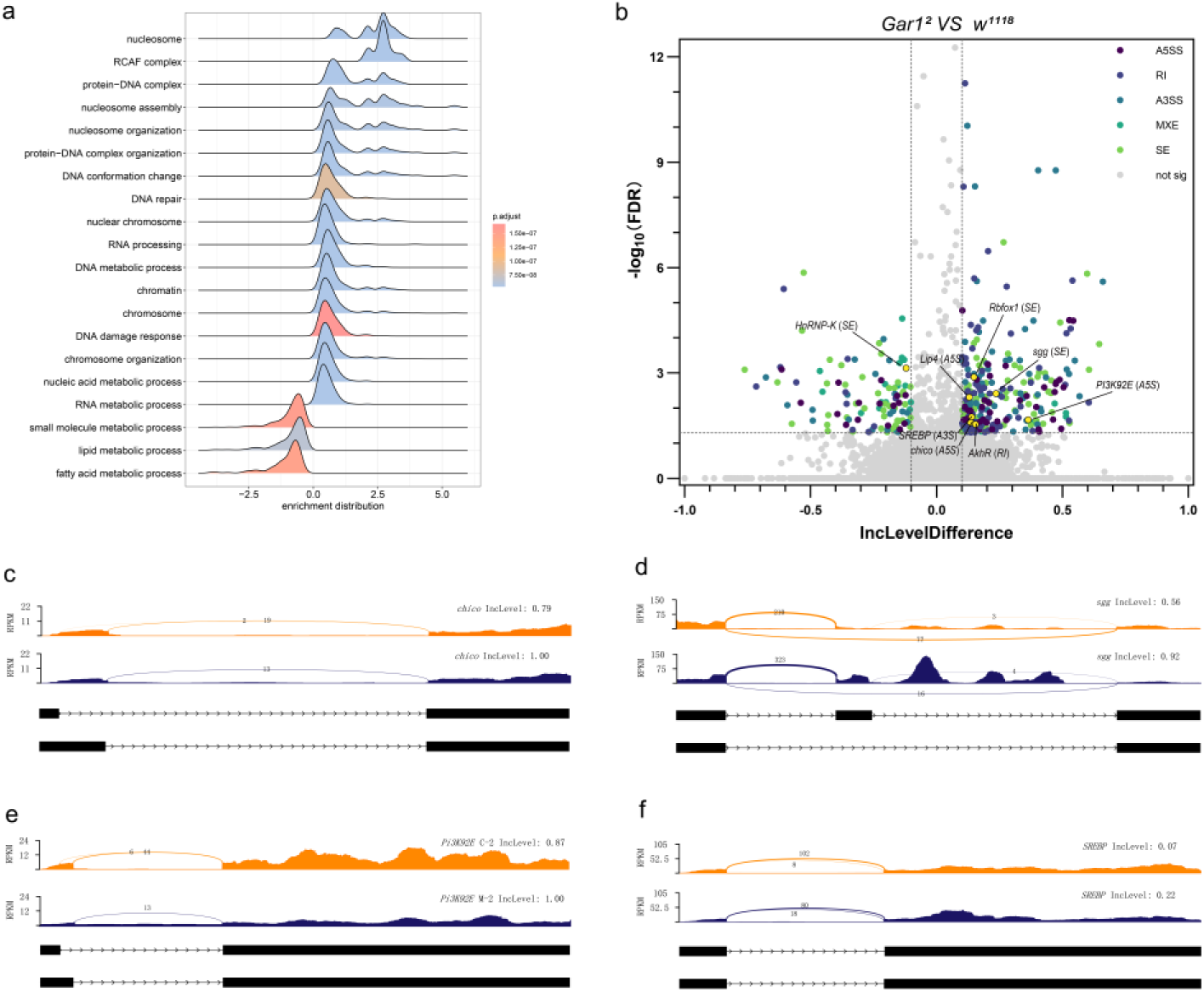
*Gar1* regulates lipid metabolism and alternative splicing. a) Positively enriched (enrich distribution > 0) and negatively enriched (enrich distribution > 0) GO terms identified by Gene Set Enrichment Analysis in *Gar1²* mutants compared to *w^1118^*controls. b) The distribution of alternative splicing events in *Gar1²* mutants relative to *w^1118^* controls. Significant splicing events, including skipped exon (SE), mutually exclusive exon (MXE), alternative 5’-Splice site (A5SS), alternative 3’-Splice site (A3SS) and retained intron (RI) were selected with a false discovery rate < 0.05 and an absolute inclusion level difference > 0.1. The highlighted circles indicated the splicing related and insulin related genes. c–f) Sashimi plots depicting splicing patterns of *chico*, *PI3K92E*, *Gsk3β (sgg)*, and *SREBP*. The inclusion levels of specific exons or alternative splice sites are indicated, revealing isoform switches in the relative genes. **ALT TEXT :** Graphs depicting that *Gar1* regulated lipid metabolism and insulin pathway at both transcriptional and post-transcriptional levels, with subfigures labelled from a to f. a) illustrate significantly upregulated and downregulated GO processes. b) Volcano plot display distribution of splicing events, with notable genes labeled. c-f) represent alternative splicing changes upon loss of *Gar1*.

To assess the functional consequences of AS changes in the insulin pathway, we examined the localization of the tGPH reporter, an *in vivo* reporter for PI3K activity (Britton et al., 2002). In the larval fat body at 68-72 h AEL, we examined the fluorescence intensity and distribution of tGPH between the membrane and cytosol. Knockdown of *Dkc1* and *Gar1* reduced the overall tGPH fluorescence by 35.96% and 47.07%, respectively (Fig. 6a and 6b). In control flies (*da-Gal4*), the tGPH signal was detected in the membrane, cytosol, and nucleus of both the salivary glands and fat bodies(Fig. 6a). Notably, *Gar1* knockdown specifically caused a more pronounced decrease in membrane-localized tGPH (Fig. 6c). Then we performed a similar analysis of tGPH in salivary gland cells. As expected, the fluorescence intensity was dramatically decreased following the knockdown of *Dkc1* or *Gar1* (Fig. 6d and 6e). Lines were drawn between the nuclei of adjacent cells to measure the distribution of the fluorescence signal (Fig. 6d; black lines). The normalized intensity readings consistently remained at high levels (greater than 80%), with a distinct peak observable at the membrane in both control (*da-Gal4*) and *da>Dkc1 RNAi* groups (arrows in Fig. 6f and 6h). In contrast, the *Gar1 RNAi* sample displayed a flat trough on the membrane with no detectable peaks (arrow in Fig. 6g). These findings indicate that the dysfunction of the box H/ACA snoRNP complex impairs insulin signaling effector activity, revealing a strong association between box H/ACA snoRNP and the insulin pathway.

**Fig. 6.**
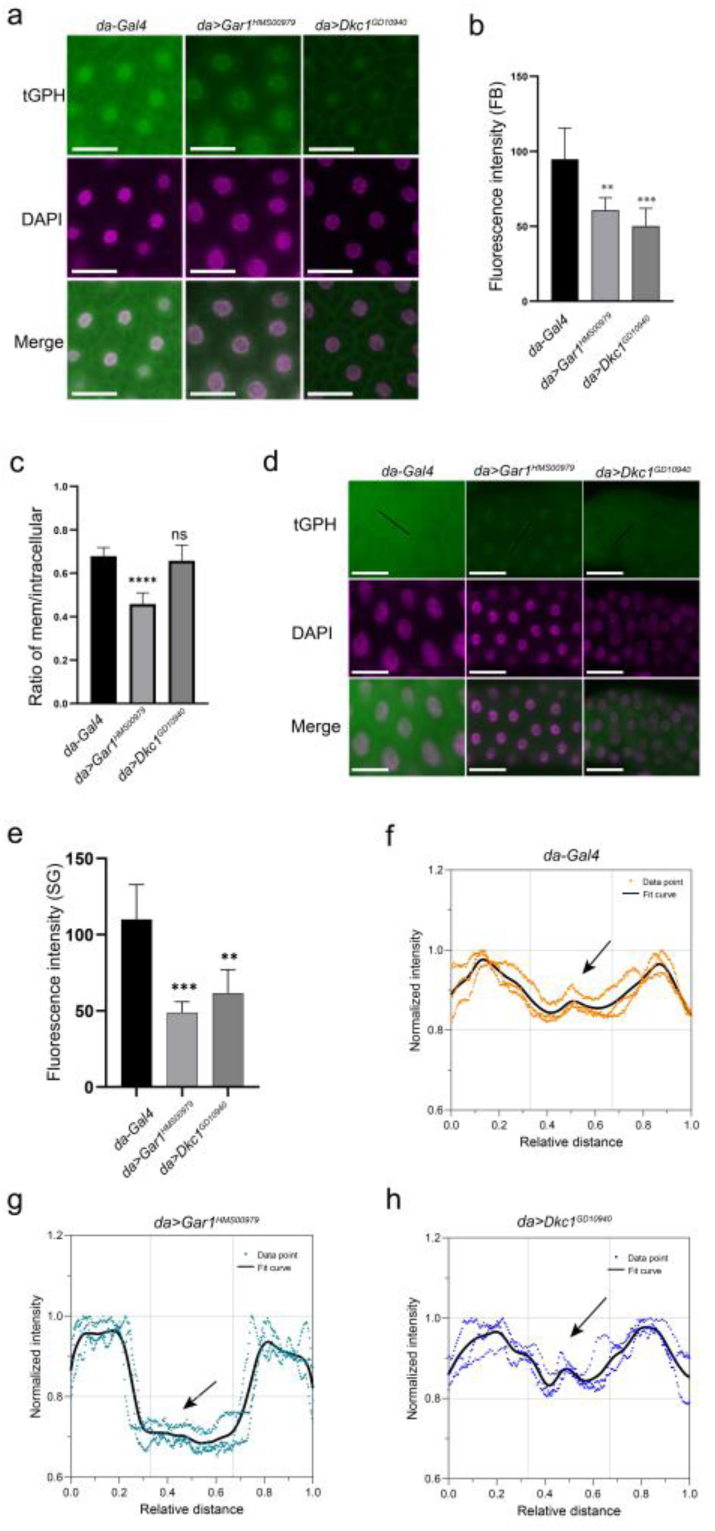
Decreased insulin signaling activity in H/ACA snoRNP dysfunction. a) Subcellular localization of the tGPH reporter (green) in third-instar larval fat body. Nuclei are stained with DAPI (Magenta). Scale bar: 50 μm. b) A significant reduction in tGPH fluorescence intensity was observed in both *Gar1* RNAi and *Dkc1* RNAi fat bodies. Statistical analysis was determined by one-way ANOVA, followed by Dunnett’s multiple comparisons test(*da-Gal4.tGPH/+* vs. da*.tGPH*, *Gar1^GD11196^*, df = 15, adjusted *p* = 0.0021; *da-Gal4.tGPH/+* vs. da*.tGPH*, *Dkc1^GD10940^*, df = 15, *p* = 0.0002). c) Alteration in tGPH fluorescence distribution in *Gar1* RNAi mutant fat bodies. Statistical analysis was determined by Brown-Forsythe ANOVA, followed by Dunnett’s T3 multiple comparisons test (*da-Gal4.tGPH/+* vs. da*.tGPH*, *Gar1^GD11196^*, df = 9.409, adjusted *p* <0.0001; *da-Gal4.tGPH/+* vs. da*.tGPH*, *Dkc1^GD10940^*, df = 7.716, adjusted *p* =0.7921). d) tGPH localization in third-instar larval salivary glands of *Gar1* and *Dkc1* RNAi mutants. Green (tGPH), Magenta (DAPI). Scale bar: 50 μm. e) The fluorescence intensity of tGPH in *Gar1* RNAi and *Dkc1* RNAi salivary glands. Statistical analysis was determined by one-way ANOVA, followed by Dunnett’s multiple comparisons test (*da-Gal4.tGPH/+* vs. da*.tGPH*, *Gar1^GD11196^*, df = 12, adjusted *p* = 0.0001; *da-Gal4.tGPH/+* vs. da*.tGPH*, *Dkc1^GD10940^*, df = 12, adjusted *p =* 0.0011). f, g, and h) Normalized fluorescence distribution in salivary gland lines. In this assay, homozygous *da-Gal4.tGPH / da-Gal4.tGPH* crossed to w^1118^, *Gar1^GD11196^* or *Dkc1^GD10940^*. The offsprings were raised in the 25 °C after synchronization. The genotypes are shown in the figures. ****, *p* < 0.0001;***, *p* < 0.001; **, *p* < 0.01; ns, not significant. **ALT TEXT**: Graph depicting the insulin activity decreased upon loss of *Gar1*, illustrated with the intensity measurement and distribution analysis of PI3K indicator. a and b) show reduced lipid droplet size in the fat bodies following knockdown of all four box snoRNP components, with corresponding statistical analyses presented in d), f) and h), respectively.

Given the previous findings that impaired box H/ACA snoRNP function disrupted lipid storage, we explored the genetic interactions in lipid deposition between *Gar1* and key genes of the insulin pathway. To this end, we employed RNAi strains targeting insulin signaling components and examined the resulting changes in larval salivary glands and fat bodies. As expected, we found that the knockdown of *lin-28* and *foxo* restored lipid deposition defects in *Gar1* mutant. In the fat body, the knockdown of *lin-28* (*lin-28^Th01982.N^*) and *foxo* (*foxo^HMS00793^*) driven by *ppl-Gal4* displayed reduced lipid droplet size compared to wild type, with average diameter of the 30 largest lipid droplets measuring 12.62 μm and 12.02 μm, respectively (Fig. 7a and 7b). Notably, *foxo* knockdown in *Gar1^HMS00979^* mutants restored the lipid droplet size to wild-type levels (Fig. 7a and 7b), indicating that FOXO and GAR1 function in the same pathway and complementarily regulate lipid storage in the adipose tissue. Conversely, *lin-28* knockdown in the *Gar1^HMS00979^* background was similar to that of *lin-28* single mutant, indicating that *lin-28* acts downstream of *Gar1.* We further examined genetic interactions in the salivary gland. Knockdown of *lin-28* or *foxo* alone using *ppl-Gal4* alone did not induce ectopic lipid droplet accumulation. Importantly, the knockdown of either gene suppressed ectopic lipid droplet formation in *Gar1^HMS00979^* mutant (Fig. 7c), confirming that *lin-28* and *foxo* function downstream of *Gar1*. Taken together, these results demonstrate that box H/ACA snoRNP complexes maintain lipid homeostasis through the insulin pathway, underscoring the tight functional connection between snoRNP and metabolic processes.

**Fig. 7.**
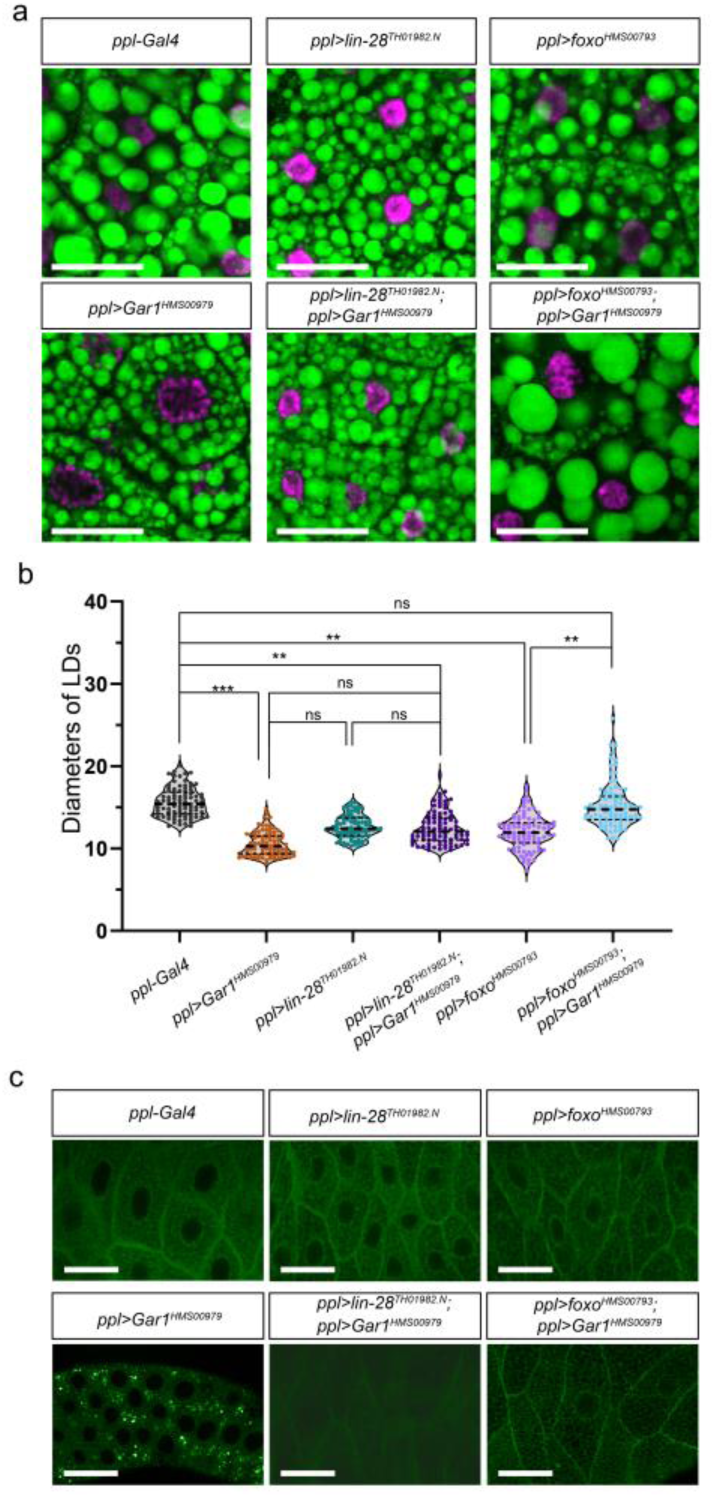
*Gar1* acts upstream of the insulin signaling pathway genetically. a) Bodipy 493/503 staining of lipid droplets in the fat body of wandering-stage larvae with the indicated genotypes. Green: Bodipy 493/503 (LDs); Magenta: DAPI (nuclei). Scale bar: 50 μm. b) Size distribution of LDs shown in (a). Statistical analysis was performed by two-way ANOVA. LD’ diameters in control and mutants. Violin plots show Max30 LDs from three figures in each genotype (n = 90 LDs per genotype). Box plots inside violins show median and quartiles. Statistical analysis was determined by one-way ANOVA using mean values, followed by Tukey’s multiple comparisons test (*ppl-Gal4* vs *ppl-Gar1^HMS00979^*, df = 12, adjusted *p* = 0.0001; *ppl- Gal4* vs *ppl>lin-28^TH01982.N^*, df = 12, adjusted *p* = 0.0119; *ppl-Gal4* vs *ppl> Gar1^HMS00979^ / lin-28^TH01982.N^*, df = 12, adjusted *p* = 0.0089; *ppl-Gal4* vs. *ppl>foxo^HMS00793^*, df = 12, adjusted *p* = 0.0029; *ppl-Gar1^HMS00979^* vs *ppl>lin-28^TH01982.N^*df = 12, adjusted *p* = 0.0979; *ppl-Gar1^HMS00979^* vs *ppl> Gar1^HMS00979^ / lin-28^TH01982.N^*, df = 12, adjusted *p* = 0.1296; *ppl>lin-28^TH01982.N^* vs *ppl> Gar1^HMS00979^ / lin-28^TH01982.N^*, df = 12, adjusted *p* >0.9999; *ppl-Gar1^HMS00979^ vs ppl> Gar1^HMS00979^ / foxo^HMS00793^,* df = 12, adjusted *p =* 0.0003*; ppl>foxo^HMS00793^* vs *ppl> Gar1^HMS00979^ / foxo^HMS00793^*, df = 12, adjusted *p* = 0.0071; *ppl-Gal4* vs. *ppl> Gar1^HMS00979^ / foxo^HMS00793^*, df = 12, adjusted *p* =0.9931). ***, *p* < 0.0001; **, *p* < 0.01; **, *p* < 0.05; ns, not significant. c) Knockdown of *lin-28* or *fox*o suppresses the ectopic lipid accumulation in the *Gar1^HMS00979^* mutants. Green: Bodipy 493/503 (LDs). Scale bar: 50 μm. **ALT TEXT**: Graph depicting the genetic interaction between insulin related genes and *Gar1*, illustrated with the morphology analysis (a, c) and statistical visualization of lipid deposition (b).

## Discussion

Adipose tissue dysfunction is a major emerging driver of metabolic syndrome, frequently characterized by excessive lipid accumulation in non-adipose tissues, such as the liver, muscle, pancreas, and renal sinus (Neeland et al., 2024). In the present study, a genetic screen was designed using the *DGAT* overexpression background. Using alterations in LD size as the phenotypic readout, we identified some well-known lipogenic and lipolytic genes (Table 1), indicating that this screen was effective. A key advantage of this screen was its capacity to distinguish fat body specific changes (Class VI and VII, Fig. 1g) from ectopic lipid deposition in the salivary glands (Class I, II, and III, Fig. 1b – 1d). Additionally, DIC is an effective technology for identifying subtle changes in lipid droplets without disrupting their native states. In fact, DIC microscopy enables the tracking of single lipid droplet (LD) dynamics over multi-day timescales (Lyn et al., 2010). In summary, this screen provides a high throughput platform for analyzing changes in both non-adipose and adipose cells in *Drosophila*.

We identified *Gar1,* which encodes a box H/ACA snoRNA-binding protein that regulates lipid storage. Knockdown of *Gar1* function led to dramatic changes in lipid deposition in both the salivary gland and fat body (Fig. 3a and S2a). The homozygous *Gar1* null mutant arrested at the L2 stage (Fig. 4i and 4j). This phenotype mimics the delayed larval development caused by *Gar1* deletion in *C. elegans*, indicating that GAR1 has a conserved function (Spaulding et al., 2022). GAR1 was initially identified in yeast for its involvement in rRNA processing and interactions with snRNAs (Girard et al., 1992) and associates with NHP2, DKC1, and NOP10 to form the snoRNP complex, regulating protein synthesis, mRNA splicing, and maintenance of genome integrity(Watkins et al., 1998; Henras et al., 1998; Mitchell et al., 1999; Pogacic et al., 2000; Kiss et al., 2010). Furthermore, we found that knockdown of *Dkc1, Nhp2* or *Nop10* led to decreased lipid storage (Fig. 3e–3h), and disruption of *Dkc1* led to lethality at the L2 stage (Fig. 4b–4e, 4l, 4m), indicating the dysfunction of other H/ACA snoRNA-binding protein phenocopies *Gar1* mutant. These results demonstrate that all four proteins function integrally, which may explain the essential roles of snoRNPs in developmental processes and energy homeostasis documented in *Drosophila*, human cells, and *Arabidopsis* (Angrisani et al., 2018; Armendariz et al., 2025; Belli et al., 2019; Li et al., 2023; Zeng et al., 2022). For example, NOP56 (component of box C/D snoRNP) and DKC1 were required for root formation and plant height development in *Arabidopsis thaliana*, with concomitant defects in pseudouridylation and 2’-O-methylation in loss of function mutants (Li et al., 2023). Another study demonstrated that downregulation of *Fib*, which encodes the box C/D component, attenuated ethanol stress and activation of the Ethanol Stress Response Element (ESRE), and the ESRE was rescued by supplementation with polyunsaturated fatty acids (Armendariz et al., 2025). We hypothesized that snoRNP dysfunction leads to impaired lipid metabolism and developmental defects by mediating key pathways that participate in lipid metabolism and development.

To verify this hypothesis, we analyzed alternative splicing changes in *Gar1^2^* mutant compared to the wild type and found that AS events occurred in key regulators of the insulin signaling pathway (Fig. 5b–5f). Insulin signaling is modulated at the post-transcriptional level by alternative splicing. For instance, the insulin receptor has two isoforms, INSR-A and INSR-B, which result from the skipping or inclusion of exon 11, respectively (Li and Huang, 2024; Malakar et al., 2016). Additionally, several genes related to glucose and lipid metabolism are regulated by AS, such as *LIPIN1, PPARγ, FADS1*, and *mTOR* (Kaminska, 2025; Liu and Klein, 2018). These results confirmed that genes in the insulin pathway were alternatively spliced, suggesting a key role for AS in metabolic regulation. *Dkc1* depletion was known to attenuate AKT-mTOR signaling through suppressed phosphorylation of 4EBP1, AKT, P70S6K, and GSK-3β (Maiello et al., 2022). Consistent with this, we also observed reduced tGPH signals in both *Dkc1-KD* and *Gar1-KD* flies (Fig. 6a–6h), indicating a functional link between the insulin signaling pathway and the snoRNP complex. *Lin-28* promotes *de novo* fatty acid synthesis by binding to the SREBP-1/SCAP mRNAs and enhancing SREBP-1 expression (Zhang et al., 2019). Insulin-dependent phosphorylation leads to nuclear exclusion of FOXO, whereas nuclear retention is linked to insulin resistance, reduced lipogenesis, and enhanced lipolysis (Lee and Dong, 2017). In present study, downregulation of *Gar1* led to reduced lipid storage and increased EFA levels. These lipid storage defects were suppressed by the knockdown of either *lin-28* or *foxo* (Fig. 7a–7c), indicating a functional connection between the box H/ACA complex and the insulin signaling pathway. However, the specific snoRNA effectors and downstream targets of the box H/ACA snoRNP complexes in lipid homeostasis remain unclear.

Several studies have highlighted the roles of snoRNAs in lipid metabolism. Box C/D snoRNAs, U32a/U33/U35a, confer resistance to lipotoxicity induced by a high-fat diet (Michel et al., 2011). The C/D snoRNA U60 knockdown resulted in impaired cholesterol trafficking (Brandis et al., 2013). H/ACA snoRNA U17 deficiency disrupts the intracellular cholesterol transport in CHO cells (Jinn et al., 2015). Furthermore, mice deficient in *U32a, U33, U34*, and *U35a* exhibited improved glucose tolerance, enhanced insulin secretion, reduced ROS production, and better responses to diabetogenic stimuli (Lee et al., 2016). In fact, the adipose tissue is far more than a passive energy storage organ. As an active endocrine entity, it engages in a complex bidirectional dialogue with the brain by releasing *upd2*, thereby systemically regulating insulin secretion and whole-body metabolic homeostasis (Ingaramo et al., 2020; Rajan and Perrimon, 2012). Furthermore, insulin secretion is modulated by *Rbfox2* via alternative splicing to maintain glucose homeostasis (Moss et al., 2023). Therefore, we hypothesized that snoRNAs, as key epigenetic regulators, may also play a profound role in this sophisticated adipose-brain crosstalk, offering a perspective for understanding the metabolic regulation network. Furthermore, identifying and characterizing specific snoRNAs involved in lipid homeostasis will provide targets for the pathological basis of metabolic syndromes.

## Data availability statement

Fly lines are available upon request. RNA-seq data are available in the NCBI Sequence Read Archive (SRA) database (accession number PRJNA1439316). https://www.ncbi.nlm.nih.gov/sra/PRJNA1439316. The authors affirm that all other data necessary for confirming the conclusions of the article are present within the article, figures, and tables. Supplemental material available at GENETICS online.

## Acknowledgments

We thank Dr. Jiahuai Han for providing the EP stocks; Dr. Pierre Leopold for providing the *ppl-Gal4* line; Dr. Renjie Jiao and Dr. Haiyang Chen for sgRNA design and transgenic plasmid construction; Dr. Jingyan Zhang for suggestions on improving the figures; and Editage (www.editage.cn) for English language editing.

